# A workflow for the detection of antibiotic residues, measurement of water chemistry and preservation of hospital sink drain samples for metagenomic sequencing

**DOI:** 10.1101/2023.09.22.558957

**Authors:** G Rodger, K Chau, P Aranega-Bou, A Roohi, G Moore, KL Hopkins, S Hopkins, AS Walker, N Stoesser

## Abstract

**Background:** Hospital sinks are environmental reservoirs that harbour healthcare-associated (HCA) pathogens. Selective pressures in sink environments, such as antibiotic residues, nutrient waste and hardness ions, may promote antibiotic resistance gene (ARG) exchange between bacteria. However, cheap and accurate sampling methods to characterise these factors are lacking.

**Aim:** To validate a workflow to detect antibiotic residues and evaluate water chemistry using dipsticks. Secondarily, to validate boric acid to preserve the taxonomic and ARG (“resistome”) composition of sink trap samples for metagenomic sequencing.

**Methods:** Antibiotic residue dipsticks were validated against serial dilutions of ampicillin, doxycycline, sulfamethoxazole and ciprofloxacin, and water chemistry dipsticks against serial dilutions of chemical calibration standards. Sink trap aspirates were used for a “real-world” pilot evaluation of dipsticks. To assess boric acid as a preservative of microbial diversity, the impact of incubation with and without boric acid at ∼22°C on metagenomic sequencing outputs was evaluated at Day 2 and Day 5 compared with baseline (Day 0).

**Findings:** The limits of detection for each antibiotic were: 3µg/L (ampicillin), 10µg/L (doxycycline), 20µg/L (sulfamethoxazole) and 8µg/L (ciprofloxacin). The best performing water chemistry dipstick correctly characterised 34/40 (85%) standards in a concentration-dependent manner. One trap sample tested positive for the presence of tetracyclines and sulfonamides. Taxonomic and resistome composition were largely maintained after storage with boric acid at ∼22°C for up to five days.

**Conclusions:** Dipsticks can be used to detect antibiotic residues and characterise water chemistry in sink trap samples. Boric acid was an effective preservative of trap sample composition, representing a low-cost alternative to cold-chain transport.

## Introduction

Antimicrobial resistance (AMR) is a significant healthcare challenge, and a global public health threat [1-4]. Hospitals represent a major site for the emergence and dissemination of multi-drug resistant (MDR) pathogens, such as carbapenemase-producing and extended-spectrum β-lactamase (ESBL)-producing *Enterobacterales* (CPE and ESBL-E, respectively), particularly in critical care units [5-7]. These MDR pathogens can colonise healthcare-associated (HCA) environmental reservoirs such as hospital sinks and contribute to healthcare-associated infections (HAIs), with many studies linking HAIs to sink drains [8-10], and successfully reducing HAI incidence via sink decontamination or removal [11, 12].

Metagenomics is increasingly used to characterise species and resistome diversity in polymicrobial samples such as those from sink drains, but the results may be significantly affected by sampling methodology, including storage conditions [13] and delays between sample collection and processing such as those resulting from transportation to the laboratory [14, 15]. Sample stabilisers, such as boric acid, which has been shown to limit bacterial overgrowth in urine samples and is widely used for community urine sampling [16], may offer a cheap and straightforward approach to maintaining sample microbial composition for environmental surveys using metagenomics.

The impact of antibiotics on bacterial and AMR gene (ARG) persistence is of particular concern in healthcare wastewater systems since these represent a confluence of patient waste, antibiotic and chemical residues, and nutrients, creating a favourable environment for ARG exchange in bacterial communities [17, 18]. For example, beta-lactam antibiotics and their metabolites are excreted in many human fluid types, potentially exerting selective pressures in disposal sites [18]. Moreover, polymicrobial biofilms in hospital sink drains demonstrate high rates of ARG transfer [6, 19, 20]. Additionally, the presence of disinfectants, such as chlorine and ethanol, non-antibiotic pharmaceuticals and heavy metal pollutants including titanium dioxide can also promote ARG transfer [20-22].

Understanding how healthcare-associated environmental drivers such as antibiotic residues and water chemistry contribute to the selection and dissemination of MDR pathogens within the hospital estate is a prerequisite to optimising infection prevention and control (IPC) and improving patient outcomes. However, rapidly and accurately evaluating these environmental drivers has been considered costly and challenging owing to the need for specialised analytical methods, and time-sensitive sample deterioration. Portable, cheap and easy-to-use dipstick approaches to measuring antibiotic concentrations and water chemistry would be of benefit.

In this study we piloted an easy-to-use workflow using dipsticks to detect the presence of antibiotic residues and evaluate water chemistry in hospital sink traps and water chemistry of hospital tap water. In addition, we validated the use of boric acid to preserve the taxonomic and resistome composition of hospital sink trap samples during transport prior to metagenomic sequencing.

## Methods

### Antibiotic residue dipstick evaluation

The QuaTest BTSQ 4-in-1 (Beta/Tetra/Sulfa/Quino) rapid test kit (Ringbio, China) was validated according to the manufacturer’s instructions using serial dilutions of ampicillin (Cambridge Bioscience, UK), doxycycline (Merck Life Science, UK), sulfamethoxazole (Insight Biotechnology Ltd, UK) and ciprofloxacin (Cambridge Bioscience, UK) from 1 mg/ml (acting as a positive control) to the limits of detection (LoD) for each antibiotic. This dipstick was chosen as it evaluates antibiotics commonly used in human healthcare settings.

### Water chemistry dipstick evaluation

Three water quality dipstick kits (Bebapanda Upgrade 14-in-1 reagent strips [China], SaySummer 16-in-1 reagent strips [China] and Qguai 9-in-1 test strips [China]) were evaluated using calibration standard dilutions of: copper, chloride, nitrate, nitrite, hardness, pH and alkalinity (Merck Life Science, UK). A fourth dipstick, Sensafe Boris’s Silvercheck strips (USA), was validated using silver calibration standard dilutions (Merck Life Science, UK).

For dipsticks that performed best on the calibration standards we conducted a “real-world” dipstick evaluation. Three hospital sink traps (A, B & C) were aspirated (50mls/trap) as described previously [23] and tested using the Bebapanda Upgrade 14-in-1 reagent strips, Sensafe Boris’s Silvercheck strips and the QuaTest BTSQ 4-in-1 test kit. Aliquots of tap water were collected from each sink after running each tap for 30 seconds, then tested using both the Bebapanda and Sensafe Silvercheck dipsticks.

### Validation of boric acid as a preservative of microbial composition in samples

Following dipstick evaluations, the three trap aspirates were also used to validate boric acid as a preservative of microbial diversity. Three timepoints were evaluated: baseline (Day 0), and at days 2 (Day 2) and 5 (Day 5) processed with and without boric acid (n=5 samples per sink). For the baseline samples DNA was extracted immediately using the PowerSoil kit (Qiagen); the day 2 +/-boric acid and day 5 +/-boric acid samples were incubated at ∼22°C for two and five days respectively prior to metagenomic DNA extraction. Metagenomes (n=15) were sequenced on the Illumina MiSeq (v3 kit, 2×300bp). Metagenome taxonomic and resistome profiles were generated with ResPipe (v.1.6.1) [24]. To address the difference in sequencing effort across samples, all samples were randomly subsampled to 475,190 reads (the minimum observed). Mean absolute percentage error (MAPE) and Bray-Curtis dissimilarity of genera were used to assess preservation of baseline taxonomy and sample-level dissimilarity respectively. Normality was confirmed using the Shapiro-Wilk test before paired sample t-testing to evaluate for statistically significant differences in taxonomic and ARG distributions (p<0.05) between samples.

## Results

### Antibiotic residues can be detected with dipsticks, with varying limits of detection (LoD)

The LoDs for the QuaTest BTSQ 4-in-1 test kit for each antibiotic were: 3 µg/L (ampicillin), 10 µg/L (doxycycline), 20 µg/L (sulfamethoxazole) and 8 µg/L (ciprofloxacin). No cross-reactivity nor false positive results were observed, and control lines were present in each test ensuring validity of each result.

### Chemistry dipsticks performed differently, with variable limits of detection

Overall, the Bebapanda Upgrade strips correctly characterised 34/40 (85%) of calibration standard concentrations whereas the SaySummer and Qguai strips were less reliable, correctly characterising only 23/40 (56%) of the biochemistry calibration standard concentrations (Table 1). Incorrect classifications were at the lower end of the concentration ranges, consistent with varying LoDs, with Bebapanda incorrectly classifying copper 1 mg/L, residual chlorine 0.5 mg/L, nitrate 10 mg/L, and nitrite 1, 5 and 10 mg/L. SaySummer and Qguai strips incorrectly classified copper 1 and 10 mg/L, residual chlorine 0.5, 1 and 3 mg/L, nitrate 10, 25 and 50 mg/L, nitrite 1, 5 and 10 mg/L, hardness 25 and 50 mg/L (plus 125 mg/L SS only), pH 7.0 (Q only), and finally alkalinity 40, 80 and 120 mg/L. Sensafe Boris’s Silvercheck strips correctly characterised all 6 silver dilutions (0, 0.05, 0.1, 0.25, 0.5 and 1.0 mg/L) .

**Table 1.**
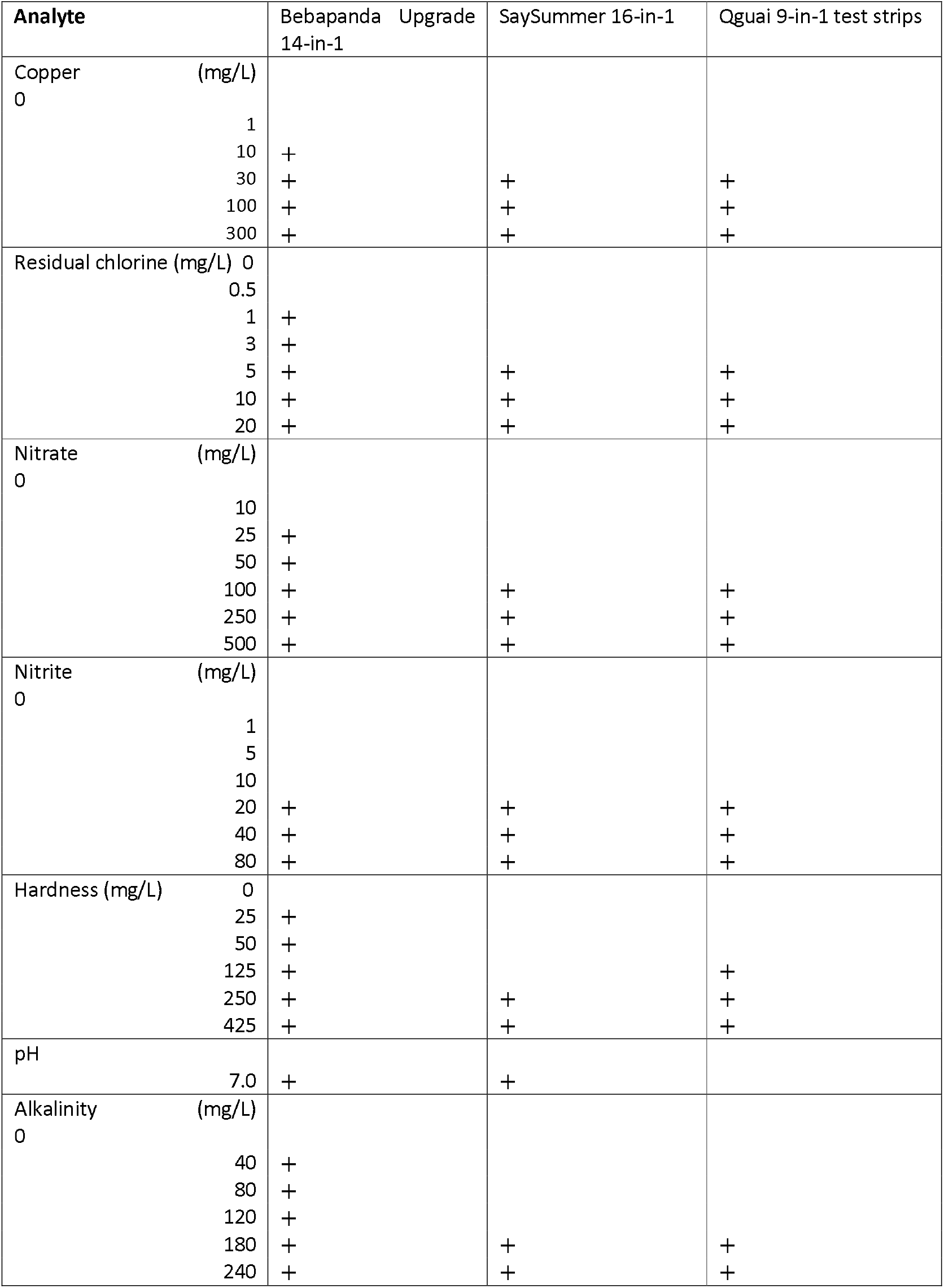
Results of the dipstick tests on calibration standards of the chemical analytes being tested. “+” denotes cases where the analyte was detected with the dipstick, a blank value cases where the analyte was not detected with the dipstick.

| Analyte | Bebapanda Upgrade<br>14-in-1 | SaySummer 16-in-1 | Qguai 9-in-1 test strips |
| --- | --- | --- | --- |
| Copper (mg/L) |  |  |  |
| 0 |  |  |  |
| 1 |  |  |  |
| 10 | + |  |  |
| 30 | + | + | + |
| 100 | + | + | + |
| 300 | + | + | + |
| Residual chlorine (mg/L) |  |  |  |
| 0 |  |  |  |
| 0.5 |  |  |  |
| 1 | + |  |  |
| 3 | + |  |  |
| 5 | + | + | + |
| 10 | + | + | + |
| 20 | + | + | + |
| Nitrate (mg/L) |  |  |  |
| 0 |  |  |  |
| 10 |  |  |  |
| 25 | + |  |  |
| 50 | + |  |  |
| 100 | + | + | + |
| 250 | + | + | + |
| 500 | + | + | + |
| Nitrite (mg/L) |  |  |  |
| 0 |  |  |  |
| 1 |  |  |  |
| 5 |  |  |  |
| 10 |  |  |  |
| 20 | + | + | + |
| 40 | + | + | + |
| 80 | + | + | + |
| Hardness (mg/L) |  |  |  |
| 0 |  |  |  |
| 25 | + |  |  |
| 50 | + |  |  |
| 125 | + |  | + |
| 250 | + | + | + |
| 425 | + | + | + |
| pH |  |  |  |
| 7.0 | + | + |  |
| Alkalinity (mg/L) |  |  |  |
| 0 |  |  |  |
| 40 | + |  |  |
| 80 | + |  |  |
| 120 | + |  |  |
| 180 | + | + | + |
| 240 | + | + | + |

### Dipsticks can be used on “real-life” sink trap and water samples

Dipsticks were easy-to-use on the “real-life” sink trap samples. The trap aspirate from sink C tested positive for the presence of tetracycline and sulfonamide antibiotic classes, denoted as the absence of a test line and/or line colour intensity less than that of the control line (Figure 1). Chemical indicators varied across sink trap aspirates and tap water samples to some extent, highlighting that measuring these parameters may provide valuable context in an epidemiological survey. Alkalinity values appropriately mirrored pH values, and one sink drain (sink C) demonstrated a low pH and the presence of bromine and chlorine, possibly consistent with the application of a cleaning agent (Table 2).

**Table 2.** Sink trap aspirate and tap water sample chemistry measurements using Bebapanda Upgrade 14 in 1 reagent dipsticks and Sensafe Boris’s Silvercheck dipsticks.

| Analyte | Sink trap A | Tap A | Sink trap B | Tap B | Sink trap C | Tap C |
| --- | --- | --- | --- | --- | --- | --- |
| Alkalinity (mg/L) | 240 | 180 | 240 | 180 | 0 | 120 |
| pH | 8.4 | 7.8 | 8.4 | 7.8 | 6.2 | 7.2 |
| Hardness (mg/L) | 125 | 125 | 125 | 125 | 425 | 425 |
| Fluoride (mg/L) | 25 | 25 | 25 | 25 | 25 | 0 |
| Sulfite (mg/L) | 0 | 0 | 0 | 0 | 0 | 0 |
| Nitrite (mg/L) | 0 | 0 | 0 | 0 | 0 | 0 |
| Nitrate (mg/L) | 0 | 25 | 25 | 25 | 0 | 25 |
| Bromine (mg/L) | 0 | 0 | 0 | 0 | 20 | 0 |
| Free chlorine (mg/L) | 0 | 0 | 0 | 0 | 5 | 0 |
| Mercury (mg/L) | 0 | 0 | 0 | 0 | 0 | 0 |
| Chromium (mg/L) | 0 | 0 | 0 | 0 | 5 | 0 |
| Iron (mg/L) | 0 | 0 | 0 | 0 | 5 | 5 |
| Copper (mg/L) | 0 | 0 | 0 | 1 | 1 | 0 |
| Lead (mg/L) | 20 | 20 | 20 | 20 | 20 | 20 |
| Silver (mg/L) | 0 | 0 | 0 | 0 | 0 | 0 |

**Figure 1.**
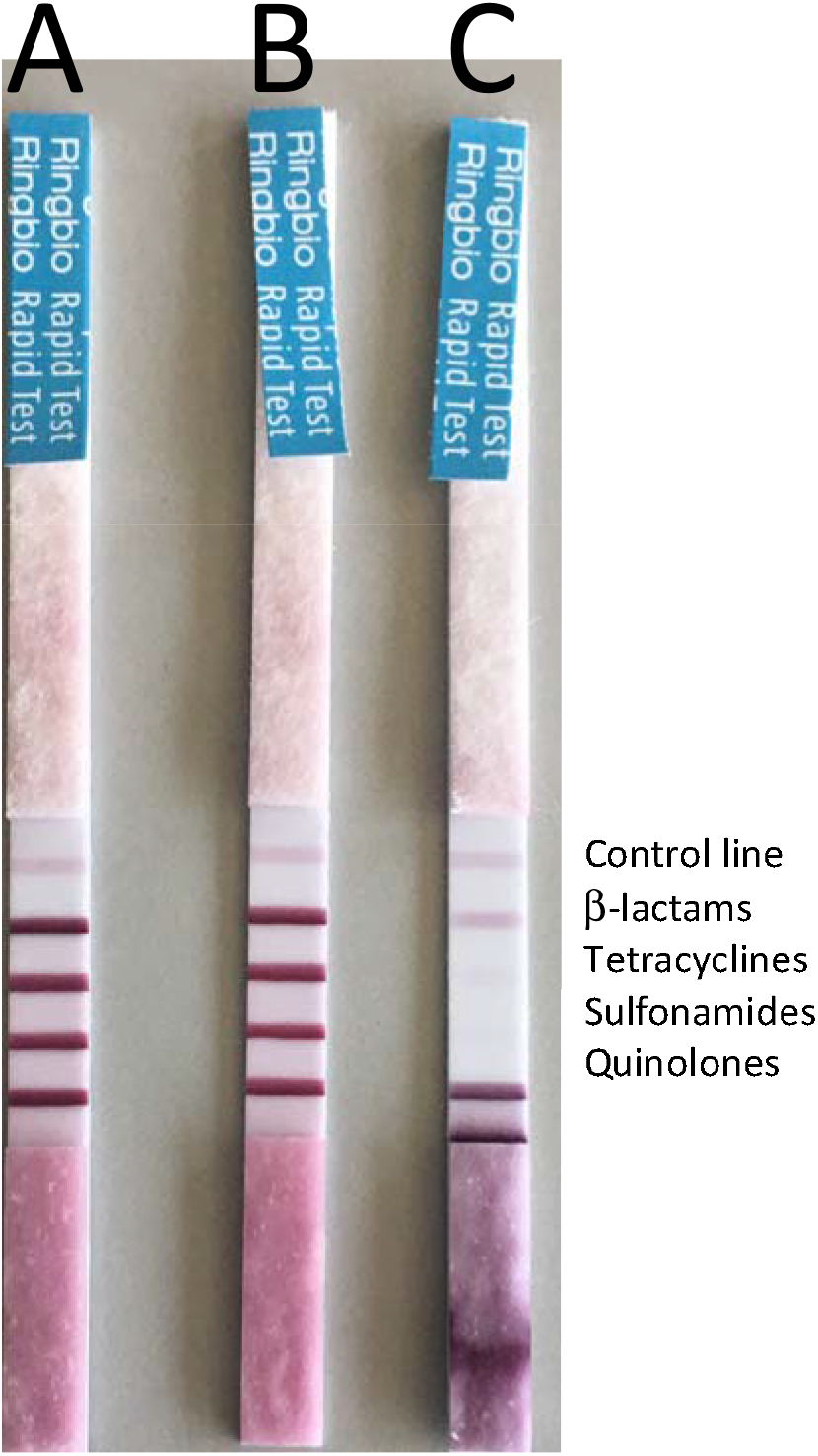
Antibiotic residues of tetracycline and sulfonamide classes of antibiotics detected in sink trap sample C. **A** control line indicates the dipstick is performing; antibiotics are characterised as present if the antibiotic-specific line is less dark than the control line or absent.

### Boric acid preserves the microbial taxonomic and AMR gene composition of sink trap samples for metagenomics

Boric acid significantly preserved the microbial composition of sink trap aspirates. Proteobacteria dominated all samples, but apparent overgrowth was prevented in samples supplemented with boric acid (Figure 2, top panel). 1-MAPE scores (with higher scores indicating greater similarity with baseline samples) indicated baseline (Day 0) taxonomic diversity and abundance were most closely preserved in samples containing boric acid, with significant divergence from baseline in sink A samples without boric acid (p=0.01) (Figure 2, bottom panel). Samples without boric acid diverged from baseline by Day 2 whereas samples with boric acid still resembled baseline at Day 5; however, decreasing 1-MAPE between days 2 and 5 was observed for all samples even with boric acid, indicating that the preservative effect of boric acid on composition is time-sensitive (Figure 2, bottom panel).

**Figure 2.**
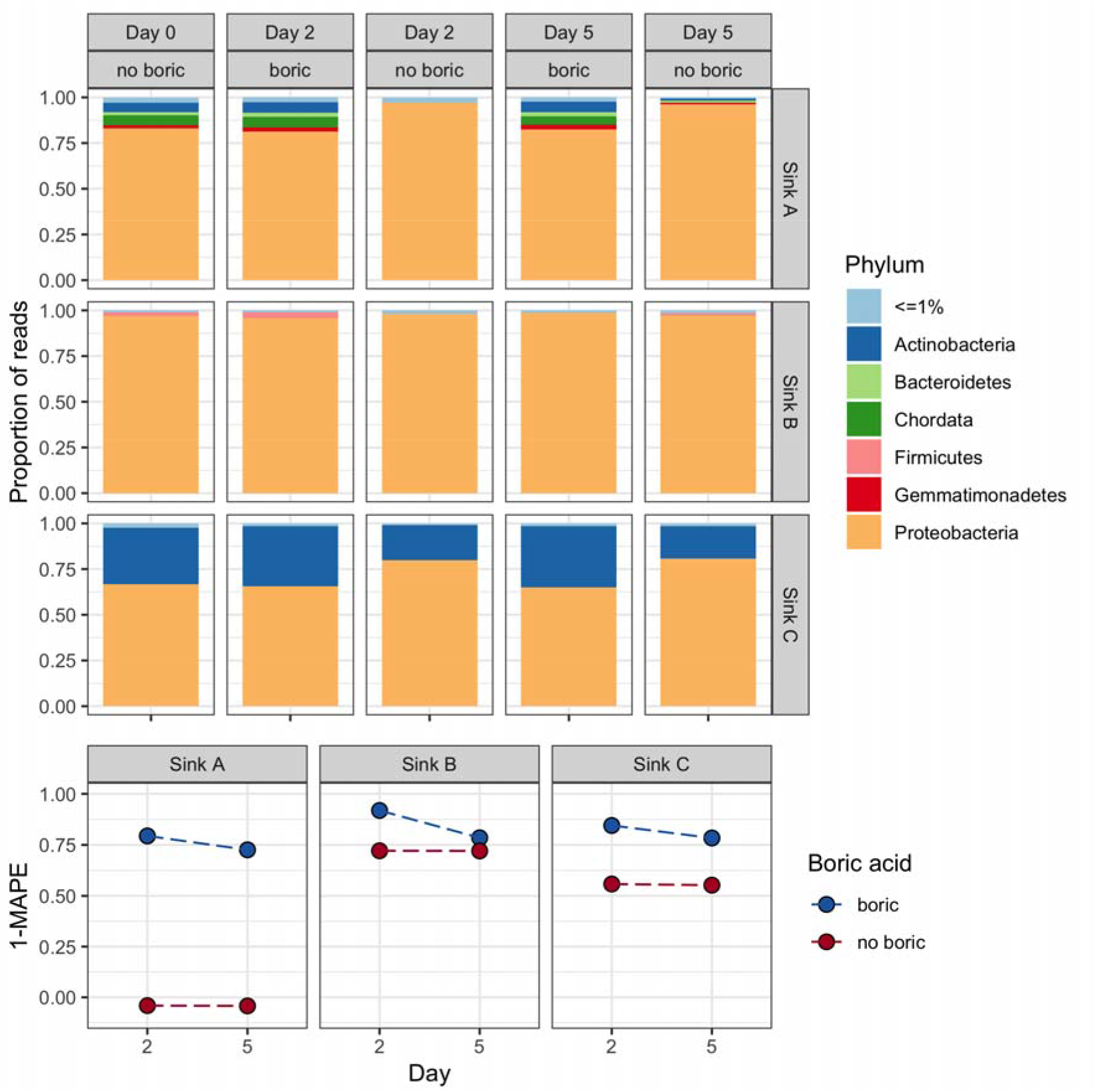
Taxonomic distributions in samples with and without boric acid. (Top panel) Relative abundances of phyla across sink samples faceted by sink (A, B, C), processing day and use of boric acid as a preservative. <= represents phyla under 0.01 proportion of reads. (Bottom panel) Mean absolute percentage error (MAPE) of taxonomic abundances between baseline (Day 0) and processing day faceted by sink and stratified by whether boric acid was used as a preservative, where 1-MAPE=1 indicates perfect preservation of baseline taxonomic distributions.

The preservative effect of boric acid was less evident for the resistome, possibly due to limited sensitivity in detecting ARGs in these samples at this sequencing depth, as demonstrated by the low numbers of sequencing reads matching AMR gene targets (Figure 3, top panel). In sink A, no ARGs were detected at baseline, making it impossible to calculate 1-MAPE scores. In sink B, where aminoglycoside resistance genes were detected at baseline, sulfonamide resistance genes were also detected by Day 5, with proportionally more divergence observed in the sample without boric acid, likely consistent with the overgrowth of a sulfonamide resistant species (Figure 3, top panel). For sink C, both samples showed a similar degree of divergence from baseline by Day 2, regardless of the presence or absence of boric acid. There was a slight increase in the 1-MAPE score for Day 5 with boric acid compared to the sample without boric acid, consistent with better preservation of ARG composition over time using boric acid. Anecdotally but of interest, the number of reads matching to ARG targets was highest for the sink C drain sample, which was also the sample in which antibiotic residues were identified. Overall, clustering patterns based on Bray-Curtis dissimilarities were observed for both taxonomy and resistome data, with distinct separation based on the sink sampled, and broadly closer clustering for cases with boric acid and baseline (Supplementary figure 1).

**Figure 3.**
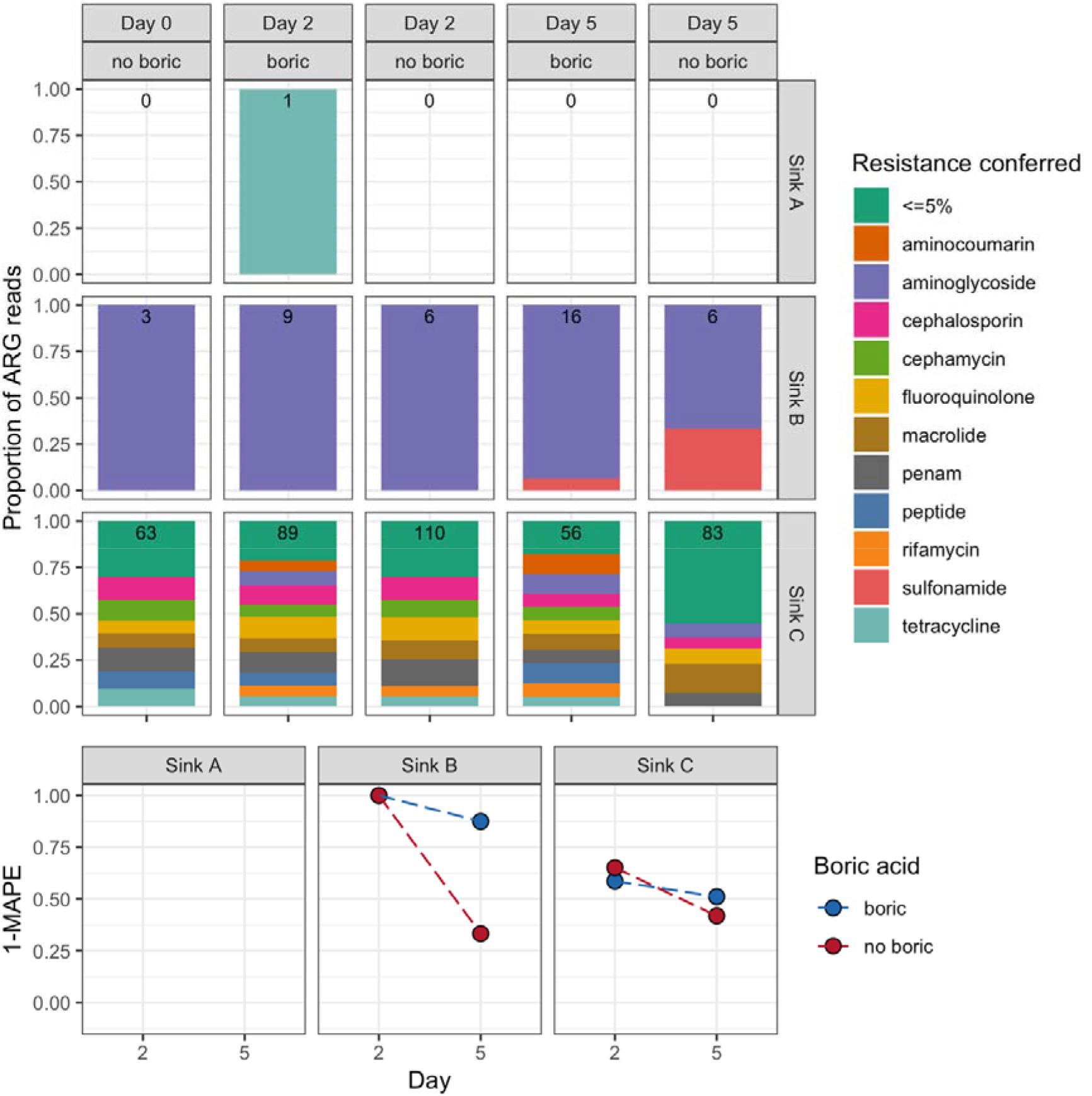
Antimicrobial resistance gene (ARG) presence in samples with and without boric acid. (Top panel) ARG reads identified in sink samples A-C by day of evaluation and with/without boric acid, represented as the proportion of reads conferring resistance to a given antimicrobial class, with the number of reads identified in each facet. <= represents resistances under 0.05 proportion of reads. (Bottom panel) Mean absolute percentage error (MAPE) of abundances between baseline (Day 0) and processing day faceted by sink sample A-C and stratified by whether boric acid was used as a preservative, where 1-MAPE=1 indicates perfect preservation of baseline ARG proportions.

## Discussion

In this pilot study we validated an easy-to-use workflow to both monitor hospital sink drains for commonly used antibiotic classes and evaluate water chemistry in sink traps and tap water, demonstrating the QuaTest BTSQ 4-in-1 antibiotic dipstick and Bebapanda Upgrade 14-in-1 and SenSafe Boris’s Silvercheck strips may be suitable for this purpose. Additionally, we validated the use of boric acid to preserve the microbial composition of hospital sink drain aspirates, mitigating transport-associated compositional changes, and facilitating more representative taxonomic and resistome profiling using metagenomics. We sampled three sinks in a healthcare setting to characterise ease of use and generate some example data.

The presence of antibiotics in hospital sink drains may facilitate persistence of AMR genes, horizontal gene transfer and emergence of MDR bacteria. Previously, Voigt *et al* [17] identified high concentrations of antibiotics from hospital sink drains, shower drains and toilets within oncology and neurological clinics, suggesting the presence of antibiotic residues in hospital wastewater corresponds to heavy antibiotic usage typical in such clinics. Effective and cheap monitoring of such environmental selection pressures could be of relevance to mitigating AMR selection in the environment. The easy-to-use QuaTest qualitative dipstick is a lateral flow immunochromatographic assay rapidly indicating antibiotic presence in a sample by the absence of a line for the corresponding antibiotic, or a line with a lower intensity in comparison to the control line. This assay identifies the presence of 40 beta-lactams, tetracyclines, sulfonamides and quinolones. Our validation against serial dilutions of known antibiotic concentrations indicates good sensitivity and specificity for all four classes of antibiotics.

Four different dipstick brands were used to measure water chemistry, including one specifically for silver concentrations. Although all dipsticks performed well in quantifying water chemistry at higher concentrations, the Bebapanda Upgrade 14-in-1 strips worked best at the lower concentrations of the eight diluted standards evaluated, and the SenSafe Boris’s Silvercheck strips accurately characterised silver concentrations. When applied to sink trap and tap water samples from three hospital sinks, some differences were noted. Piped water is generally hard in our region (∼260-280 mg/L) [25] as it contains large amounts of magnesium and calcium, but testing in our sinks demonstrated results lower (sinks A, B) and higher than this (sink C). Sinks A and B could perhaps be supplied from a different water source. For sink C, it may be that the tap from this particular sink was infrequently cleaned and a mineral plaque inside the tap was contributing to the hardness value sampled. Fluoride is not added to the water supply in this region but occurs naturally in drinking water supplies, as reflected in the results, whereas the water is chloraminated or chlorinated [25], but we were only able to detect free chlorine in trap aspirate from sink C which may have been from a cleaning product. The presence of lead is perhaps from lead service pipes or an inaccuracy in the dipstick colour chart given this analyte was not evaluated as part of our standard dilutions. Bromine, an oxidant which can be used as a sanitiser or disinfectant, was only detected in the trap aspirate of sink C which may again have been residual post-cleaning.

Biologically active environmental samples such as sink trap aspirates may undergo compositional changes during transport, potentially impacting the accuracy of subsequent analyses and interpretation. Factors such as temperature fluctuation, nutrient depletion, incubation time, residual disinfectants or cleaning products and handling practices can lead to shifts in microbial populations and DNA degradation. Although these issues can be predominately mitigated through immediate deep freezing and cold-chain transport practices, these approaches incur significant cost and present a logistical challenge. Our study demonstrates that boric acid is an effective short-term preservative of the taxonomic composition and probably resistome of sink trap samples for 2-5 days at room temperature, although our observations on preservation of the resistome were less clear, probably driven by the relatively low numbers of reads mapping to ARG targets at the sequencing depth used. Interestingly, most samples clustered by the sampled sink regardless of boric acid status for both taxonomic and resistome ordinations, suggesting sink-level differences (e.g. location, usage, design) are stronger drivers of variation than composition shifts occurring over five days storage. However, this observation needs to be considered with caution given the small numbers of sink traps evaluated.

In addition to the small number of sink trap/tap samples analysed, there were several limitations to this study. We assessed dilutions of standards for only 8 of the 14 chemicals evaluated on the chemistry dipsticks, as standards for the other analytes were not readily available. Dilution was undertaken in sterile purified water rather than a matrix mimicking a sink trap or tap water sample, as these matrices would be difficult to simulate and using real-life samples means that background concentrations of these chemicals may already be present. For the antibiotic dipstick a downside is that it gives a binary, qualitative result that does not differentiate between antibiotics within each of the four classes or any indication of the detection of multiple antibiotics within each class.

## Conclusions

This study evaluated and piloted easy-to-use dipsticks to detect the presence of four classes of antibiotic residues and measure water chemistry parameters from sink trap aspirates and tap water. Taxonomic and resistome composition of hospital trap samples were largely maintained after storage with boric acid at ∼22°C for up to five days, facilitating the collection of environmental samples for multi-site studies. Performing simple dipstick tests to detect antibiotic residues and measure water chemistry parameters in hospital wastewater and accurately evaluating pathogen and AMR gene burden in sink reservoirs may contribute to strategies to monitor and mitigate the impact of AMR selection in hospital environments.

