## Supplementary Figure 1 for "A workflow for the detection of antibiotic residues, measurement of water chemistry and preservation of hospital sink drain samples for metagenomic sequencing"

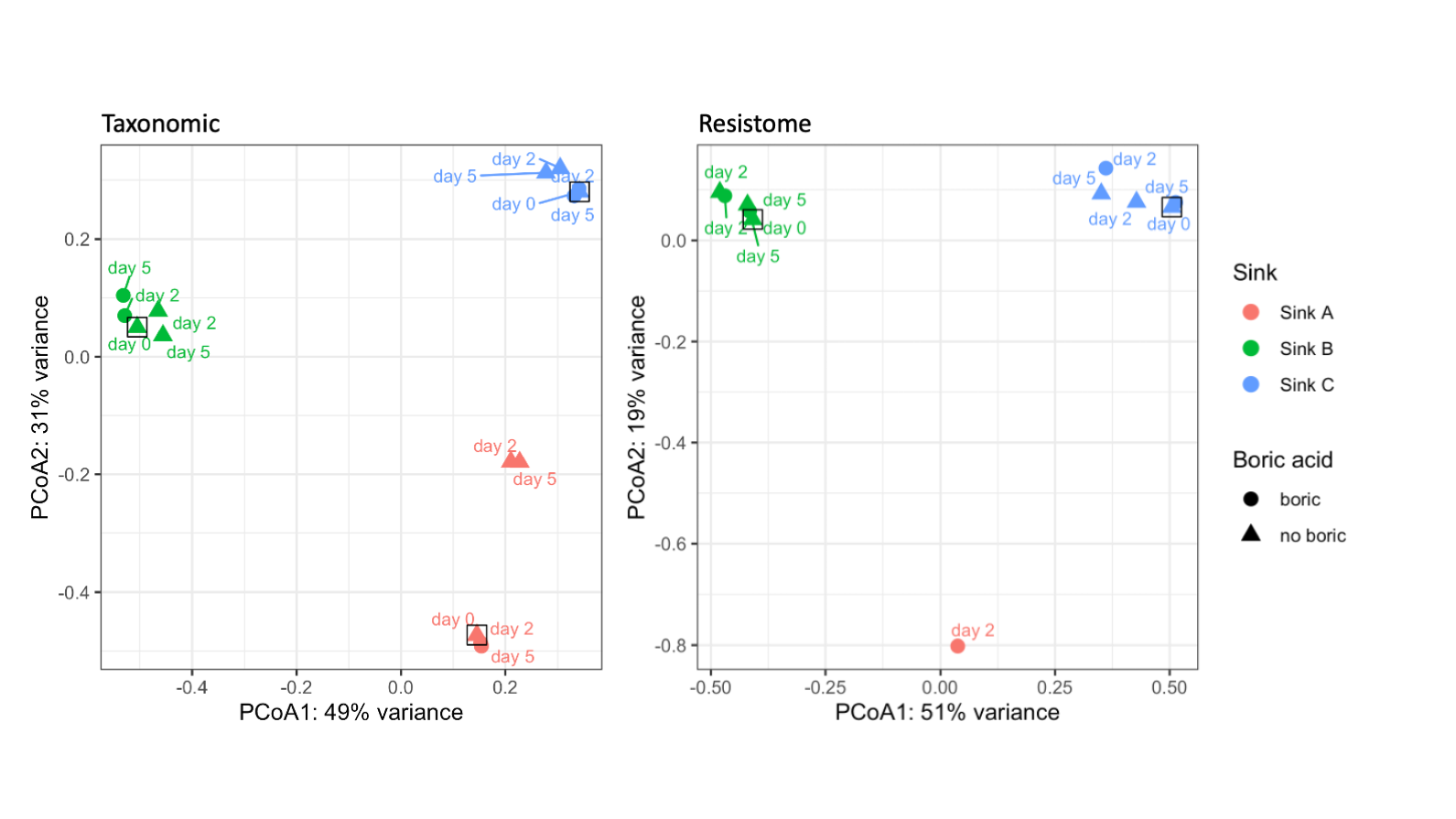

**Supplementary figure 1 Principal coordinate analyses of sample-level Bray-Curtis dissimilarities for (left panel) taxonomic and (right panel) resistome content.** Points represent individual samples with colour denoting sink and shape denoting the use of boric acid, with baseline samples highlighted (square outline). N.B. no AMR genes were detected for Sink A Day 0 or 5 +/- boric acid.
